# SARS-CoV-2 Omicron subvariants evolved to promote further escape from MHC-I recognition

**DOI:** 10.1101/2022.05.04.490614

**Authors:** Miyu Moriyama, Carolina Lucas, Valter Silva Monteiro, Yale SARS-CoV-2 Genomic Surveillance Initiative, Akiko Iwasaki

## Abstract

SARS-CoV-2 variants of concern (VOCs) possess mutations that confer resistance to neutralizing antibodies within the Spike protein and are associated with breakthrough infection and reinfection. By contrast, less is known about the escape from CD8^+^ T cell-mediated immunity by VOC. Here, we demonstrated that all SARS-CoV-2 VOCs possess the ability to suppress MHC I expression. We identified several viral genes that contribute to the suppression of MHC I expression. Notably, MHC-I upregulation was strongly inhibited after SARS-CoV-2 infection *in vivo*. While earlier VOCs possess similar capacity as the ancestral strain to suppress MHC I, Omicron subvariants exhibit a greater ability to suppress surface MHC-I expressions. Collectively, our data suggest that, in addition to escape from neutralizing antibodies, the success of Omicron subvariants to cause breakthrough infection and reinfection may in part be due to its optimized evasion from T cell recognition.

**Significance:** Numerous pathogenic viruses have developed strategies to evade host CD8^+^ T cell-mediated clearance. Here, we demonstrated that SARS-CoV-2 encodes multiple viral factors that can modulate MHC-I expression in the host cells. We found that MHC-I upregulation was strongly suppressed during SARS-CoV-2 infection *in vivo*. Notably, the Omicron subvariants showed an enhanced ability to suppress MHC-I compared to the original strain and the earlier SARS-CoV-2 variants of concern (VOCs). Our results point to the inherently strong ability of SARS-CoV-2 to hinder MHC-I expression and demonstrated that Omicron subvariants have evolved an even more optimized capacity to evade CD8 T cell recognition.

## Introduction

SARS-CoV-2 has continued to evolve since it was first detected in Wuhan, China in December 2019. Beginning in late 2020, waves of SARS-CoV-2 variants of concern (VOCs) with increased transmissibility and immune evasion capacity have emerged. Increasing breakthrough infection and reinfection events are associated with the emergence of VOCs (1, 2). Breakthrough infections and reinfections are likely driven by significant increases in transmissibility (3), evasion from innate immunity (4, 5), and escape from neutralization by vaccine/infection-induced antibodies (6–9). By contrast, minimal evasion of T cell epitopes has been reported for VOCs (10). In November 2021, the Omicron variant, the newest VOC declared by WHO to date, had emerged. Omicron variant then quickly outcompeted the previously dominant Delta variant and led to the largest surge in COVID-19 cases worldwide. The outstanding features of the Omicron variant are the considerably enhanced escape from the antibody neutralization (9, 11) and increased infectivity (12) than the earlier VOCs, due to its heavily mutated Spike protein. Although the Omicron variant and its subvariants harbor a far greater number of mutations in its genome compared to those in previous VOCs, T cell epitopes remain generally intact (13).

CD8^+^ cytotoxic T lymphocyte (CTL) recognizes and kills infected cells and eliminates the source of replicating viruses. Antigen presentation by major histocompatibility complex class I (MHC-I) is a critical step for the activation of antigen-specific CD8^+^ T cells and the subsequent killing of infected cells. Viral peptides processed by the cellular proteasome complex are loaded on MHC-I molecule in the endoplasmic reticulum and translocate to the cell surface to be recognized by antigen-specific CD8^+^ T cells. To successfully establish infection and replicate in the host, many viruses have acquired the ability to inhibit MHC-I processing and presentation of viral antigens (14). Likewise, SARS-CoV-2 utilizes its viral proteins to interfere with the MHC-I pathway (15–19). SARS-CoV-2 ORF8 protein induces autophagic degradation of MHC-I and confers resistance to CTL surveillance (15). Studies from the first 3 months of the pandemic showed a rapid evolution of the SARS-CoV-2 ORF8 gene including isolates with 382nt deletion spanning ORF7b-ORF8 gene region (20, 21), which is associated with robust T cell response and milder clinical outcome (22, 23). These findings collectively raised a question of whether VOC and its ORF8 protein have evolved to further enhance the ability to shut down MHC-I, thereby evading from antigen-specific memory CD8^+^ T cells established by previous infection or vaccination.

Here, we performed a systematic analysis of the capacity of SARS-CoV-2 variants to downregulate MHC-I presentation. Our data demonstrated vigorous suppression of MHC-I surface expression by the ancestral SARS-CoV-2 and minimal evolution in modulating MHC-I pathway by earlier VOCs. Remarkably, the latest Omicron subvariants have acquired an enhanced ability in modulating MHC-I pathway.

## Results

### Pre-Omicron SARS-CoV-2 variants retain similar MHC-I evasion capacity

To investigate the impact of SARS-CoV-2 infection on MHC-I expression, we infected Calu-3 cells, a commonly used human lung epithelial cell line, with SARS-CoV-2 variants and the ancestral strain (USA-WA1). We tested four variants of concern (B.1.1.7/Alpha, B.1.351/Beta, P.1/Gamma, and B.1.617.2/Delta) and three variants of interest (B.1.427/Epsilon, B.1.429/Epsilon, and B.1.526/Iota). To assess MHC-I expression levels, the cells were pre-gated for single cells and live cells (Fig.S1). Infection with SARS-CoV-2 variants reduced the viability of the cells by ~30% compared to the mock-infected condition (Fig.S1A-B). Within the live cell population, SARS-CoV-2 variants and the ancestral strain similarly downregulated MHC-I levels after infection (Fig.1A). We next examined transcriptional levels of MHC-I genes after infection with SARS-CoV-2 variants. Transcriptional levels of MHC-I genes differed depending on the variants (Fig.1B). The ancestral strain significantly downregulated HLA-A, B, and C genes as previously reported (16). B.1.1.7 and B.1.351 showed a similar reduction in HLA-A, B, and C mRNA expression as the ancestral strain. Other variants showed a weaker downregulation (B.1.526), no significant change (B.1.429), or upregulation (P.1) of HLA class I genes within the infected cells. These results indicated that SARS-CoV-2 variants maintain a similar capacity to reduce HLA transcription as the ancestral virus, except for the P.1 and B.1.429 variants. Given that P.1-infected and B.1.429-infected cells still expressed low levels of MHC-I on the surface, other mechanisms involved in the MHC-I processing and presentation pathway may be dampened by the infection leading to reduced surface MHC-I expression.

**Figure 1.**
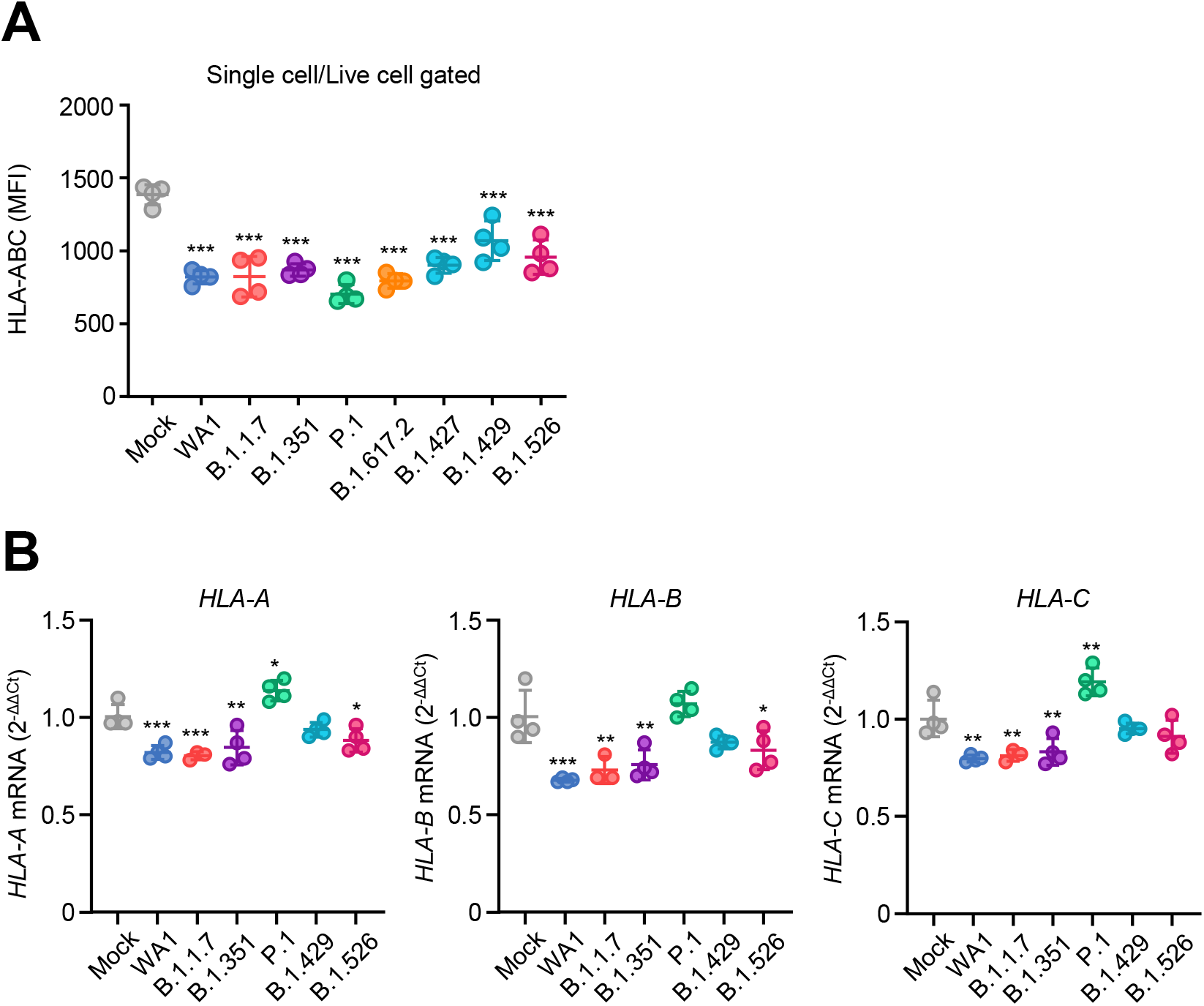
MHC-I evasion by SARS-CoV-2 variants. (A) Calu-3 cells were infected with SARS-CoV-2 variants at MOI 0.3 for 40h. The cell surface expression of HLA-ABC was analyzed by FACS. (B) Calu-3 cells were infected with SARS-CoV-2 variants at MOI 0.01 for 48h. The mRNA levels of HLA-A, B, and C were measured by RT-qPCR. Data are mean ± s.d. Data are representative of two to three independent experiments. *, p<0.05; **, p< 0.01; ***, p< 0.001

### Variant-specific mutations are found in the ORF8 gene of SARS-CoV-2

Because previous studies revealed the association between 382 nt deletion spanning ORF7b-ORF8 gene region and robust IFN-ɤ and T cell responses (22, 23), we next addressed the role of ORF8 in the differential MHC-I regulation by VOCs. We performed multiple sequence alignments of ORF8 amino acid sequences from SARS-CoV-2 variants to see if there are any non-synonymous mutations. In total, 7 non-synonymous mutations and 2 deletions were detected from 16 variants examined (Fig. S2). Notably, a premature stop codon was introduced at Q27 of B.1.1.7, which truncates the ORF8 polypeptide length and likely alters the protein functionality. The downstream mutations (R52I and Y73C) probably have no further impact on B.1.1.7 ORF8 protein because of the early translation termination by Q27stop mutation. Although pre-Omicron variants of concern (B.1.1.7/Alpha, B.1.351/Beta, P.1/Gamma, and B.1.617.2/Delta) harbored mutations or deletions in ORF8 protein, ORF8 sequence from BA.1/Omicron variant and its descendants unexpectedly remained intact. Two of the former variants of interest, B.1.429/Epsilon and B.1.526/Iota harbored V100L and T11I mutation respectively. None of the mutations and deletions were conserved among different lineages. To investigate the prevalence of mutations found in variants, we downloaded 3,059 SARS-CoV-2 genome sequence data from GISAID database (https://www.gisaid.org/). We found that the mutation in a particular amino acid is only exclusively seen in a single lineage (Fig.2A). ORF8 L84S mutation, which was detected within the first 2 months of pandemic (24) and corresponding to clade S, was not observed in any of the variants. We also observed the mutations discovered by multiple sequence alignment are generally highly prevalent, and the proportions ranged from 12.5 to 100% of the lineage (Fig.2B). These results indicate that the variant-specific mutations were acquired independently during SARS-CoV-2 evolution.

**Figure 2.**
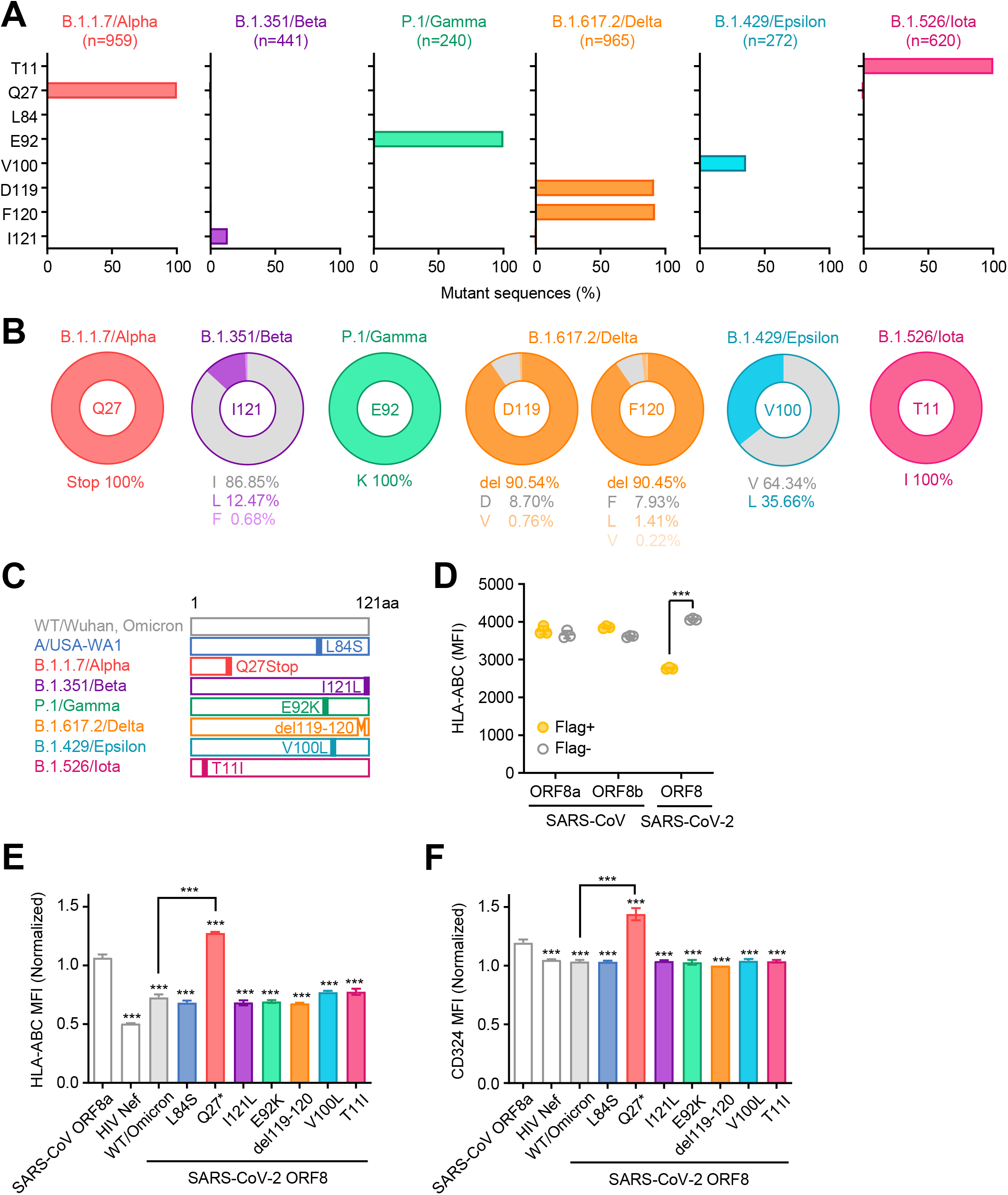
Unique mutations are found in ORF8 gene of SARS-CoV-2 variants. (A) Mutant proportion in the ORF8 genes of the indicated SARS-CoV-2 variants. The amino acid positions shown are selected based on the results of multiple sequence alignment performed in Fig.S2. The number of sequences analyzed for each lineage is shown above each graph. (B) Frequency of amino acids at the positions enriched for mutants in each variant. The amino acids shown in gray color correspond to WT. (C) Schematic diagram of ORF8 proteins from SARS-CoV-2 variants. (D-F) HEK293T cells were transfected with plasmids encoding C-terminally Flag-tagged SARS-CoV ORF8a/b, HIV Nef, SARS-CoV-2 ORF8 WT, or SARS-CoV-2 ORF8 variants. Forty-eight hours after transfection, cells were collected and analyzed for the cell surface HLA-ABC expression (D-E) and CD324 (F). Data are shown in raw MFI (D) or as the ratio of MFI in Flag+ cells to Flag-cells (E) (n=3). Data are mean ± s.d. Data are representative of two to three independent experiments. ***, p< 0.001

### Impaired MHC-I downregulation by B.1.1.7 ORF8 protein

We next tested whether variant-specific mutations alter MHC-I downregulating capability of ORF8 protein. To this end, we generated expression plasmids encoding seven ORF8 mutants from SARS-CoV-2 variants (Fig.2C), and subsequently transfected HEK293T cells with these plasmids for the detection of its effect on the surface MHC-I expression levels. We included SARS-CoV ORF8a/b proteins as negative controls, as they have been shown not to affect MHC-I expression levels (15). Since ORF8 induces degradation of MHC-I via autophagy by interacting with MHC-I and localizing in LC3-positive puncta (15), ORF8 presumably acts on MHC-I downregulation in cell-intrinsic manner. Indeed, surface MHC-I levels of the cells expressing WT ORF8 protein were much lower than those of the cells without ORF8 expression (Fig.2D). In addition to the surface MHC-I, intracellular MHC-I molecules were also decreased specifically fin ORF8-expressing cells (Fig.S3A), further supporting the direct role of ORF8 in the control of cellular MHC-I levels. The cysteine 20 residue of SARS-CoV-2 ORF8 is known to form intermolecular disulfide bonds between two ORF8 molecules and stabilize the homodimer (25, 26). Disruption of the Cys20 residue, however, did not affect the regulation of cell surface MHC-I levels (Fig.S3B). Among seven ORF8 mutants tested, six mutants including L84S, I121L, E92K, del119-120, V100L, and T11I downregulated surface MHC-I levels of the cells expressing those proteins to a similar extent to WT ORF8 protein (Fig.2E), while maintaining the expression of an irrelevant cell surface molecule, CD324 (Fig.2F). On the other hand, Q27Stop ORF8 mutant had a completely abrogated MHC-I downregulation capability compared to the WT ORF8 protein (Fig.2E). These results indicated that none of the variant-specific mutations enhanced the ability of ORF8 protein to downregulate MHC-I, and the ORF8 encoded by the B.1.1.7 lineage lost its ability to reduce surface MHC I expression.

### Multiple SARS-CoV-2 viral proteins play redundant roles in the downregulation of MHC-I

Given that B.1.1.7 and P.1 variants were able to reduce MHC-I expression levels even though these lineages retain functionally defective ORF8 mutant or are less effective in reducing HLA-I mRNA levels, we investigated the possibility that SARS-CoV-2 encodes multiple viral genes that redundantly act to suppress MHC-I expression. We generated expression plasmids encoding SARS-CoV-2 E, M, ORF7a, and ORF7b, and assessed the effect on the surface MHC-I and MHC-II expression levels of HEK293T cells following transfection with these plasmids. We also included Human Immunodeficiency Virus (HIV) Nef as a positive control for downregulating both MHC-I and MHC-II (27, 28), and SARS-CoV ORF8a/b proteins as a negative control. As expected, HIV Nef protein downregulated both MHC-I and MHC-II levels, whereas SARS-CoV-2 ORF8 specifically targeted MHC-I (Fig.3A-B). We found that in addition to ORF8, SARS-CoV-2 E, M, and ORF7a substantially downregulated MHC-I within the cells expressing these viral proteins (Fig.3C). Significant reduction of surface MHC-II levels was also observed by expression of these viral proteins (Fig.3C), albeit to a lesser extent (~20%). These results suggested that SARS-CoV-2 encodes multiple viral genes that are redundantly downregulating MHC-I likely to ensure viral evasion from MHC-I-mediated CTL recognition.

**Figure 3.**
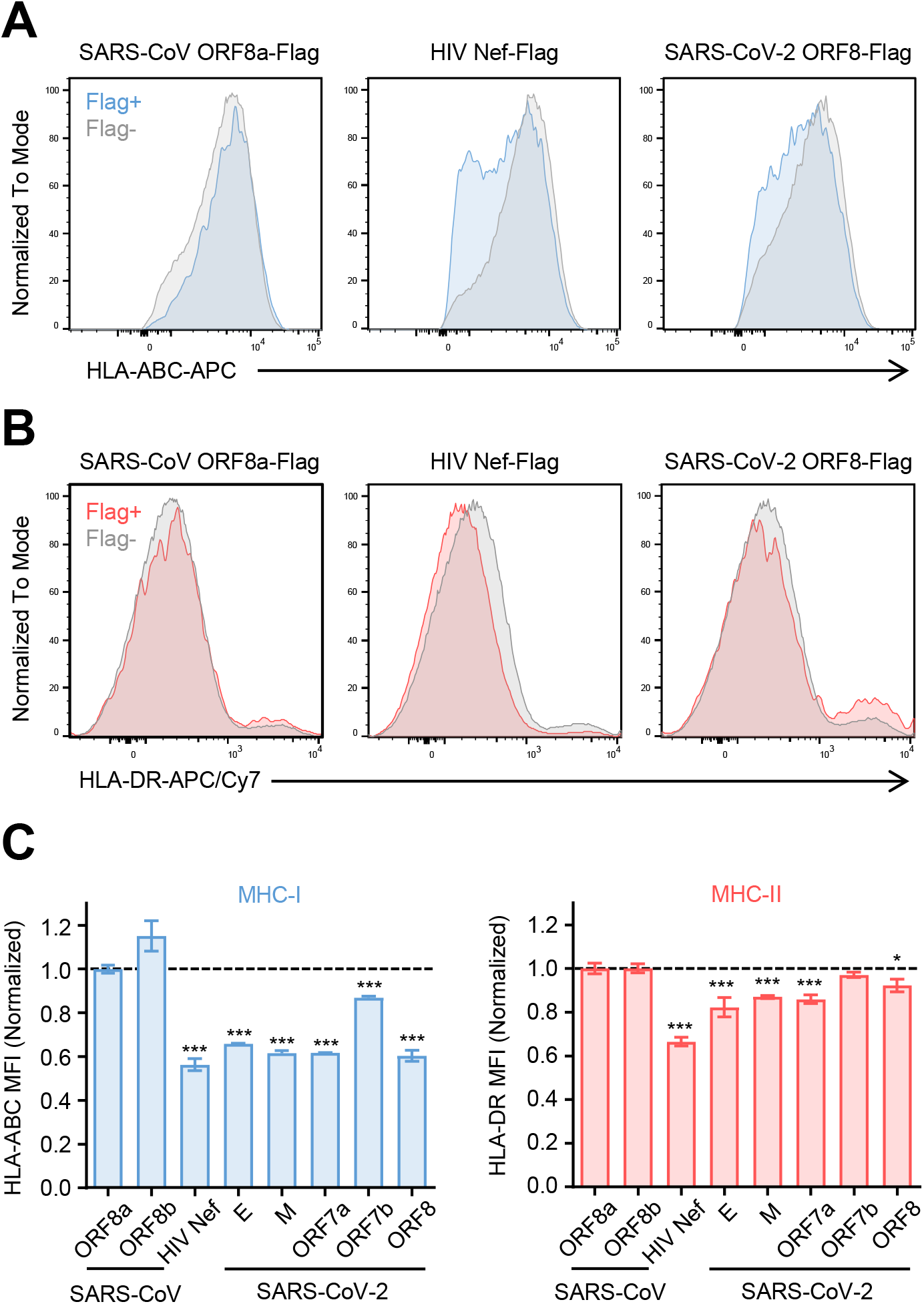
SARS-CoV-2 viral genes redundantly downregulate MHC-I. (A-C) HEK293T cells were transfected with expression plasmids encoding C-terminal Flag-tagged viral proteins as indicated. After 48 hours, surface expressions of HLA-ABC and HLA-DR were analyzed by flow cytometry. The representative histogram of HLA-ABC (A) and HLA-DR (B) of SARS-CoV ORF8a-Flag, HIV-Nef-Flag, or SARS-CoV-2 ORF8-Flag +/- cells are shown. (C) The normalized ratio of HLA surface expression in Flag+ cells to Flag-cells are shown (n=3). Data are mean ± s.d. Data are representative of three independent experiments. Statistical significance is calculated versus SARS-CoV ORF8a *, p< 0.05; ***, p< 0.001

### Superior MHC-I evasion by SARS-CoV-2 compared to influenza A virus *in vivo*

In the experiments above, we have shown that SARS-CoV-2 encodes multiple viral proteins that are targeting MHC-I expression, which can synergistically strengthen the capability of the virus to avoid MHC-I presentation. Moreover, we confirmed the previous finding that the MHC-I downregulation is a newly acquired function of SARS-CoV-2 ORF8 protein, which was not seen in SARS-CoV ORF8a/b proteins. Considering these results, we hypothesized that even the ancestral strain of SARS-CoV-2 possesses a superior MHC-I evasion strategy than other respiratory viruses. To assess this hypothesis, we infected C57BL/6J mice intranasally with a mouse-adapted strain of SARS-CoV-2 (SARS-CoV-2 MA10) or influenza A/PR8 virus and analyzed the MHC-I expression levels of lung epithelial cells at 2 days after infection. SARS-CoV-2 MA10 virus harbors 2 mutations in the Spike protein, 3 mutations in the ORF1ab, and an F7S mutation in ORF6 compared to the ancestral virus (29). Strikingly, influenza A virus induced robust upregulation of MHC-I in both infected (NP+) and uninfected (NP-) lung epithelial cells, whereas SARS-CoV-2 MA10 upregulated MHC-I only in uninfected cells (S-), and to a lesser extent than the influenza virus (Fig.4A-C). Importantly, MHC-I upregulation was completely blocked in SARS-CoV-2 MA10 infected (S+) lung epithelial cells, suggesting that SARS-CoV-2 viral proteins are strongly inhibiting MHC-I upregulation in a cell-intrinsic manner. These results indicated that SARS-CoV-2 possesses a near complete ability to shut down MHC-I induction within infected cells in vivo, a feature not found in influenza A virus.

**Figure 4.**
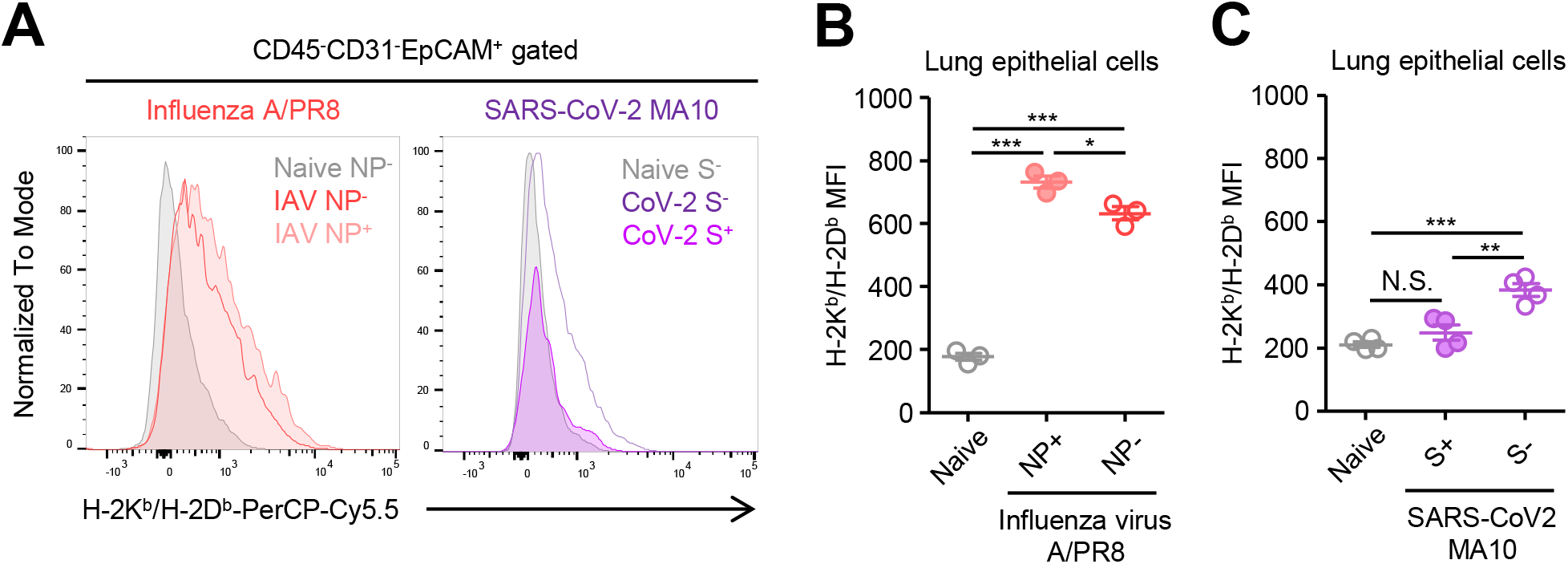
Robust suppression of MHC-I upregulation by SARS-CoV-2 *in vivo*. (A-C) C57BL/6J mice were infected intranasally with 10^5^ PFU of SARS-CoV-2 MA10 or Influenza virus A/PR8. Two days later, lungs were collected and analyzed for surface MHC-I expressions on epithelial cells of viral protein (SARS-CoV-2 Spike (S) or Influenza A virus Nucleoprotein (NP)) positive and negative populations. (n=3). The representative histograms (A) and MFI (B and C) are shown. Data are mean ± s.e.m. Data are representative of two independent experiments. *, p<0.05; **, p< 0.01; ***, p< 0.001

### Omicron subvariant downregulates MHC-I more efficiently than older isolates

Finally, we tested the ability of the more recent Omicron subvariants to counteract the MHC-I expression. We generated an A549 cell line, a human adenocarcinoma cell line that stably expresses ACE2 (A549-hACE2 cells). A549-hACE2 cells were infected with five Omicron subvariants (BA.1, BA.2.12.1, XAF, BA.4, and BA.5) along with older isolates (WA1 and B.1.429) for comparison. We further distinguished viral infected (S+) cells from bystander cells (S-) to assess the direct impact of infection on the cell surface MHC-I expression (Fig.5A), by looking at cell surface HLA-ABC raw MFI (Fig.5B-C) or normalized MFI in S+ infected cells (Fig.5D-E). Consistent with the observation in Calu-3 cells (Fig.1A), SARS-CoV-2 infection suppressed MHC-I expression in A549-hACE2 cells (Fig.5B, D). Furthermore, MHC-I reduction was specifically seen in S+ infected cells and varied between different viral strains (Fig 5B). Whereas an irrelevant cell surface marker CD324 (E-cadherin) remained unchanged upon infection with SARS-CoV-2 (Fig.5C, E). Remarkably, many of the omicron subvariants, such as BA.1, BA.2.12.1, XAF, and BA.4, had a superior capacity to reduce surface MHC-I levels compared to the older VOC (Fig.5D). These results underscore the universal capacity of all SARS-CoV-2 strains to mediate the cell-intrinsic reduction of MHC-I expression within the infected cells, and highlight the evolution of the Omicron subvariants in acquiring a superior MHC-I evasion capacity.

**Figure 5.**
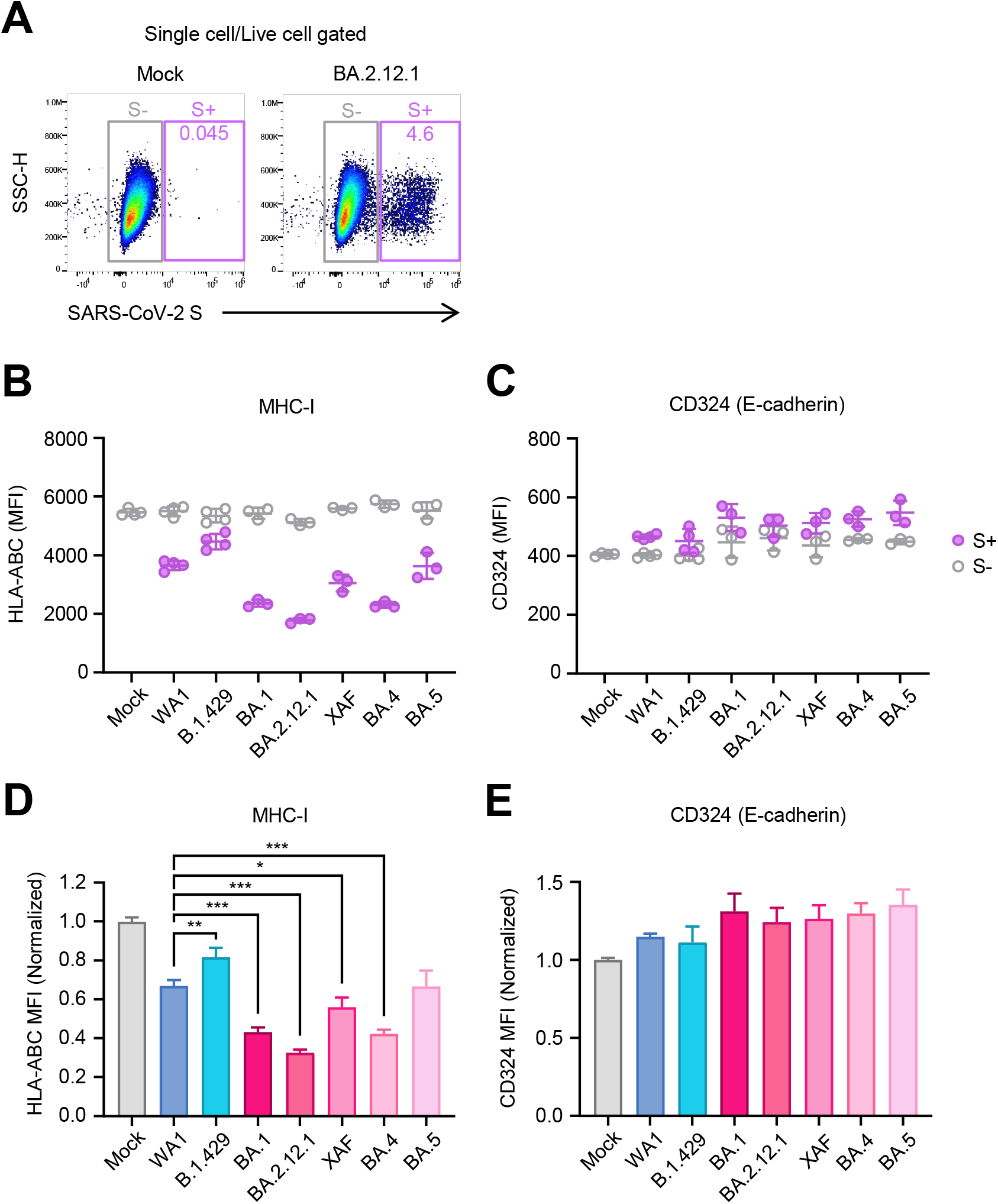
Increased suppression of MHC-I by Omicron sublineages. (A-E) A549-hACE2 cells were infected with SARS-CoV-2 variants at MOI 0.3 for 44h. Cells were collected and analyzed for surface MHC-I expressions on single live cells of SARS-CoV-2 Spike (S) positive and negative populations by FACS. (A) Representative FACS plot. Data are shown as the raw MFI (B-C) or the normalized ratio of MFI in SARS-CoV-2 S+ cells to MFI in mock cells (D-E). Data are mean ± s.d. Data are representative of two independent experiments. *, p<0.05; **, p< 0.01; ***, p< 0.001

## Discussion

CD8^+^ T cell-mediated elimination of infected cells plays an important role in the antiviral adaptive immune response. Thus, many viruses have developed ways to avoid the efficient MHC-I mediated antigen presentation to CD8^+^ T cells. In the current study, we uncovered the intrinsically potent ability of SARS-CoV-2 to shut down the host MHC-I system by using live, authentic SARS-CoV-2 variants as well as the functional analysis of variant-specific mutations in ORF8 gene, a key viral protein for both MHC-I evasion and adaptation to the host. We further identified multiple other viral genes that confer redundant function in MHC I suppression. We show that respiratory epithelial cells infected *in vivo* with SARS-CoV-2 failed to upregulate MHC-I, whereas those infected with influenza virus robustly elevated MHC-I expression. Finally, our data revealed that the most recent Omicron subvariants have superior capacity to suppress MHC-I compared to the earlier isolates.

We demonstrated that most variants of concern/interest possess unique mutations within ORF8 gene. However, none of these ORF8 mutations led to further reduction of MHC-I in cells expressing these molecules. Notably, the Omicron variant and its descendants lacked in non-synonymous mutations in their ORF8 gene, indicating that the mechanisms for superior MHC-I suppression by the Omicron sub-lineages must be located outside the ORF8 protein.

Our data showed that the ORF8 mutation in the B.1.1.7/Alpha lineage abrogated its function in MHC I modulation. Given the truncation mutation likely rendering ORF8 in the Alpha variant non-functional, this raises a question as to how such mutations might be tolerated. Multiple functions beyond MHC-I downregulation are documented for SARS-CoV-2 ORF8, which include inhibition of type I IFN, ISGs, or NF-kB signaling (30–32), epigenetic modulation through histone mimicry (33) and induction of proinflammatory cytokines from macrophages and monocytes via IL-17RA (34–37). Interestingly, several studies showed that SARS-CoV-2 ORF8 is actively secreted into the cell culture media in a signal peptide-dependent manner when it is overexpressed *in vitro* (26, 34, 38). Furthermore, ORF8 peptides and anti-ORF8 antibodies can be detected abundantly in serum of patients, suggesting the relevance of the active secretion of ORF8 to actual infection in humans (38). The Alpha variant likely acquired compensatory mechanisms that enabled its successful transmission until the next variant came along.

ORF8 is implicated in adaptation to the human host during the SARS-CoV outbreak (39, Chinese SARS Molecular Epidemiology 40), and it is known that ORF8 is the hypervariable genomic region among the SARS-CoV and bat SARS-related CoVs (41, 42). Likewise, studies from early in the COVID-19 pandemic observed the variability and rapid evolution of SARS-CoV-2 ORF8 gene (20, 43). Notably, SARS-CoV-2 isolates with 382nt deletion spanning ORF7b-ORF8 gene region were observed in Singapore (21), which correlated with robust T cell response and mild clinical outcome (22, 23). Mutations in ORF8 gene thus may play a key role in modulating viral pathogenesis and adaptation to the host by regulating MHC-I levels and ISGs.

The enhanced immune evasion by VOCs has been well documented for escape from neutralizing antibodies (6–8) and from innate immune responses (4, 5). Here we demonstrated that the ability to reduce MHC-I expression remained unchanged throughout the pre-Omicron VOC evolution. These findings suggested three important perspectives on the MHC-I evasion strategy of SARS-CoV-2. First, SARS-CoV-2 utilizes multiple redundant strategies to suppress MHC-I expression. For example, considering B.1.1.7 retained an intact ability to shut down MHC-I, the impaired MHC-I evasion by B.1.1.7 ORF8 is likely compensated by the redundant and/or compensatory functions of other viral proteins including E, M, and ORF7a. In addition, B.1.1.7 lineage has been shown to express an increased subgenomic RNA and protein abundance of ORF6 (4), which suppresses MHC-I at the transcriptional level by interfering with STAT1-IRF1-NLRC5 axis (16). The multi-tiered MHC-I evasion mechanisms thus work redundantly to ensure escape from CTL killing.

Second, MHC-I downregulation may not only impair CTL recognition of infected cells for killing but may also impair priming of CD8 T cells. Indeed, the frequency of circulating SARS-CoV-2 specific memory CD8^+^ T cells in SARS-CoV-2 infected individuals are ~10 fold lower than for influenza or Epstein-Barr virus-specific T cell populations (44), which indicates the suboptimal induction of memory CD8^+^ T cells following SARS-CoV-2 infection in human.

Third, given that the variants of concern had not further evolved to downregulate MHC-I more strongly than the original strain except for the Omicron subvariants, SARS-CoV-2 ancestral virus was already fully equipped to escape from CD8^+^ T cell-mediated immunity with respect to downregulation of MHC-I expression and is under less evolutionary pressure to further optimize the evasion strategy than those from type I IFNs or antibodies. However, mutations and evasion from particular H LA-restricted CTL epitopes have been observed in circulating SARS-CoV-2 and VOCs (45, 46). Genome-wide screening of epitopes suggested the CD8^+^ T and CD4^+^ T cell epitopes are broadly distributed throughout SARS-CoV-2 genome (47, 48), and the estimated numbers of epitopes per individual are at least 17 for CD8^+^ T and 19 for CD4^+^ T cells, respectively (48), and thus functional T cell evasion by VOCs is very limited (10). This in turn suggests that MHC-I downregulation may be a more efficient way for viruses to avoid CTL surveillance than introducing mutations in epitopes. The importance of MHC-I evasion by SARS-CoV-2 is also highlighted by the fact that no genetic mutations or variations in the MHC-I pathway has thus far been identified as a risk factor for severe COVID (COVID-19 Host Genetics 49), unlike innate immune pathways involving TLRs and type I IFNs (50).

SARS-CoV-2 infection in both human and preclinical models have shown to induce antigen-specific CD8^+^ T cell responses (51, 52), and the early CTL response correlated with a milder disease outcome in human (53). Adoptive transfer of serum or IgG from convalescent animals alone, however, is enough to reduce viral load in recipients after SARS-CoV-2 challenge in mice and non-human primates (54, 55) and neutralizing antibody is shown to be a strong correlate of protection (54, 56, 57). The protective roles of CD8^+^ T cell-mediated immunity appear to be more important in the absence of the optimal humoral responses/neutralizing antibody (54, 58). Blood anti-ORF8 antibodies can be used as the highly sensitive clinical marker for SARS-CoV-2 infection early (~14 days) after symptom onset (38, 59), which suggests the role of ORF8 in the very early stage of the disease. ORF8-mediated MHC-I downregulation can therefore precede antigen presentation and hinder priming of viral antigen-specific CD8^+^ T cell immune responses. Robust MHC-I shutdown by SARS-CoV-2 may explain in part the less effective protection by CD8^+^ T cells and the less impact of CD8^+^ T cell absence compared with humoral immunity (54).

Collectively, our data shed light on the intrinsically potent ability of SARS-CoV-2 to avoid the MHC-I mediated antigen presentation to CD8^+^ T cells. Importantly, we observed a complete inhibition of MHC-I upregulation in lung epithelial cells infected with SARS-CoV-2 at the early stage of infection in a mouse model. Since the ability of ORF8 to downregulate MHC-I is a newly acquired feature in SARS-CoV-2 ORF8 and is absent in SARS-CoV ORF8 (15), it is possible that ORF8 played a role in the efficient replication and transmission of SARS-CoV-2 in human and contributed to its pandemic potential. Our work provides insights into SARS-CoV-2 pathogenesis and evolution and predicts difficulty for CD8 T cell-based therapeutic approaches to COVID-19.

## Materials and methods

### Mice

Six to ten-week-old male C57BL6 mice were purchased from the Jackson Laboratory. All animal experiments in this study complied with federal and institutional policies of the Yale Animal Care and Use Committee.

### Cell lines and viruses

HEK293T cells and A549-hACE2 cells were maintained in DMEM supplemented with 1% Penicillin-Streptomycin and 10% heat-inactivated FBS. Calu-3 cells were maintained in MEM supplemented with 1% Penicillin-Streptomycin and 10% heat-inactivated FBS. ACE2-TMPRSS2-VeroE6 cells were maintained in DMEM supplemented with 1% sodium pyruvate, 1% Penicillin-Streptomycin and 10% heat-inactivated FBS at 37C. Influenza virus A/Puerto Rico/8/34 was kindly provided by Dr. Hideki Hasegawa (National Institute of Infectious Diseases in Japan). Virus stocks were propagated in allantoic cavities from 10-to 11-day-old fertile chicken eggs for 2 days at 35C. Viral titers were determined by standard plaque assay procedure. SARS-CoV-2 MA10 (29) was kindly provided by Dr. Ralph S. Baric (University of North Carolina at Chapel Hill). SARS-CoV-2 lineage A (USA-WA1/2020) and B.1.351b (hCoV-19/South Africa/KRISP-K005325/2020) were obtained from BEI resources. Pre-Omicron lineages (B.1.1.7(GenBank Accession: MZ202178), B.1.351a (GenBank Accession: MZ202314), P.1 (GenBank Accession: MZ202306), B.1.617.2 (GenBank Accession: MZ468047), B.1.427 (GenBank Accession: MZ467318), B.1.429 (GenBank Accession: MZ467319), and B.1.526 (GenBank Accession: MZ467323)) and Omicron sub-lineages (BA.1 (GenBank Accession: ON425981), BA.2.12.1 (GenBank Accession: ON411581), XAF (GenBank Accession: OP031604), BA.4 (GenBank Accession: ON773234), and BA.5.2.1 (GenBank Accession: OP031606)) were isolated and sequenced as part of the Yale Genomic Surveillance Initiative’s weekly surveillance program in Connecticut, United States, as previously described (60). Virus stocks were propagated and titered as previously described (8, 61). Briefly, TMPRSS2-VeroE6 cells were infected at multiplicity of infection of 0.01 for 3 days and the cell-free supernatant was collected and used as working stocks. All experiments using live SARS-CoV-2 were performed in a biosafety level 3 laboratory with approval from the Yale Environmental Health and Safety office.

### Viral genome sequence analysis

Pre-Omicron SARS-CoV-2 variant genome sequences (3,067 sequences) were downloaded from GISAID database (https://www.gisaid.org/) as of February 23, 2022. Sequences of Wuhan Hu-1 (GenBank accession: NC_045512.2) and USA-WA1/2020 (GenBank accession: MW811435.1) were obtained from NCBI Virus SARS-CoV-2 Data Hub (https://www.ncbi.nlm.nih.gov/labs/virus/vssi/#/sars-cov-2). To investigate the prevalence of amino acid mutations, we downloaded up to 965 sequences of each lineage and aligned the ORF8 nucleotide sequences using Jalview software (http://www.jalview.org/) (Waterhouse et al. Bioinformatics. 2009) by MUSCLE algorithm (Edgar RC. Nucleic Acid Res. 2004). Sequences containing undetermined nucleotides within the codon of interest were removed for analysis. ORF8 amino acid sequence alignment was conducted by Jalview software using MUSCLE algorithm.

### Viral infection

Mice were fully anesthetized by intraperitoneal injection of ketamine and xylazine, and intranasally inoculated with 50ul of PBS containing 1×10^5^ PFU of influenza virus A/Puerto Rico/8/34. For SARS-CoV-2 infection in the animal biosafety level 3 facility, mice were anesthetized by 30% v/v isoflurane diluted in propylene glycol, and 50ul of 1×10^5^ PFU of SARS-CoV-2 MA10 in PBS was intranasally delivered. For cell culture infection, cells were washed with PBS and infected with SARS-CoV-2 at a multiplicity of infection of 0.01 or 0.3 for 1 h at 37C. After 1 h incubation, cells were supplemented with complete media and cultured until sample harvest.

### Plasmids

pDONR207-SARS-CoV-2 E (#141273), pDONR207-SARS-CoV-2 M (#141274), pDONR207-SARS-CoV-2 ORF7a (#141276), pDONR223-SARS-CoV-2 ORF7b (#141277), pDONR223-SARS-CoV-2 ORF8 (#141278) were purchased from addgene (62) and used as templates for construction of plasmids expressing SARS-CoV-2 viral proteins. For HIV Nef expressing plasmid construction, NL4-3-dE-EGFP (kindly provided by Dr. Ya-Chi Ho) was used as a template. The full-length viral genes were amplified by PCR using iProof™ High-Fidelity DNA Polymerase (Bio-Rad), with templates described above and specific primers containing XhoI (XbaI for HIV Nef) and BamHI sites at the 5’ and 3’ ends, respectively. Following restriction enzyme digestion, PCR fragments were cloned into c-Flag pcDNA3 vector (addgene, #20011). For construction of plasmids expressing SARS-CoV viral proteins, oligonucleotides corresponding to both strands of SARS-CoV Tor2 (GenBank accession: NC_004718.3) ORF8a and ORF8b containing XhoI and BamHI sites at the 5’ and 3’ ends were synthesized (IDT) and cloned into XhoI-BamHI site of c-Flag pcDNA3 vector. Mutant SARS-CoV-2 ORF8 expressing plasmids were generated by standard PCR-based mutagenesis method. Integrity of inserts was verified by sequencing (Yale Keck DNA sequencing facility).

### Lung cell isolation

Lungs were harvested and processed as previously described (54). In Brief, lungs were minced with scissors and digested in RPMI1640 media containing 1mg/ml collagenase A, 30ug/ml DNase I at 37C for 45 min. Digested lungs were then filtered through 70um cell strainer and treated with ACK buffer for 2 min. After washing with PBS, cells were resuspended in PBS with 1% FBS.

### Flow cytometry

Cells were blocked with Human BD Fc Block (Fc1.3216, 1:100, BD Biosciences) in the presence of Live/Dead Fixable Aqua (Thermo Fisher) for 15 min at room temperature. Staining antibodies were added and incubated for 20 min at room temperature. Cells were washed with 2mM EDTA-PBS and resuspended in 100ul 2% PFA for 1 hour at room temperature. For intracellular staining, PFA-fixed cells were washed and permeabilized with eBioscience FoxP3/Transcription Factor Staining Buffer (Thermo Fisher) for 10min at 4C. Cells were washed once and stained in the same permeabilization buffer containing staining antibodies. After 30min incubation at 4C, cells were washed and resuspended in PBS with 1% FBS for analysis on Attune NxT (Thermo Fisher). FlowJo software (Tree Star) was used for the data analysis. Staining antibodies are as follows (Hu Fc Block Pure Fc1.3216 (BD, Cat# 564220), APC anti-HLA-ABC (Thermofisher, Cat# 17-9983-42), APC/Cy7 anti-HLA-DR (BioLegend, Cat# 307618), BV421 anti-mouse/human CD324 (Biolegend, Cat# 147319), PE anti-DYKDDDDK Tag (BioLegend, Cat# 637309), AF488 anti-SARS-CoV-2 Spike S1 Subunit (R&D Systems,Cat# FAB105403G), FITC anti-Influenza A NP (Thermofisher, Cat# MA1-7322), PE anti-mouse CD45 (BioLegend, Cat# 109808), BV421 anti-mouse CD31 (BioLegend, Cat# 102423), APC anti-mouse EpCAM (BioLegend, Cat# 118213), PerCP/Cy5.5 anti-H-2Kb/H-2Db (BioLegend,Cat# 114620)).

### Quantitative PCR

SARS-CoV-2 infected cells were washed with PBS and lysed with TRIzol reagent (Invitrogen). Total RNA was extracted using RNeasy mini kit (QIAGEN) and reverse transcribed into cDNA using iScript cDNA synthesis kit (Bio-Rad). RT-PCR was performed by CFX96 Touch Real-Time PCR detection system (Bio-Rad) using iTaq SYBR premix (Bio-Rad) and following primers (5’-3’): HLA-A (Forward: AAAAGGAGGGAGTTACACTCAGG, Reverse: GCTGTGAGGGACAC ATCAGAG), HLA-B (Forward: CTACCCTGCGGAGATCA, Reverse: ACAGCCA GGCCAGCAACA), HLA-C (Forward: CACACCTCTCCTTTGTGACTTCAA, Reverse: CCACCTCCTCACATTATGCTAACA), human GAPDH (Forward: CAACGGATTTGGTCGTATT, Reverse: GATGGCAACAATATCCACTT).

## Statistical analysis

Statistical significance was tested using one-way analysis of variance (ANOVA) with Tukey’s multiple comparison test. P-values of <0.05 were considered statistically significant.

## Acknowledgements

We thank Melissa Linehan and Huiping Dong for technical and logistical assistance. We thank Ralph Baric for kindly providing SARS-CoV-2 MA10. We thank Ya-Chi Ho for kindly providing NL4-3-dE-EGFP. We thank Craig Wilen for his technical expertise. We thank Benjamin Israelow and Tianyang Mao for critical reading of the manuscript. We also give special recognition to the services of Ben Fontes and the Yale EH&S Department for their ongoing assistance in safely conducting biosafety level 3 research. This work was in part supported by the Fast Grant from Emergent Ventures at the Mercatus Center and 1R01AI157488. A.I. is an Investigator of the Howard Hughes Medical Institute. M.M. is supported by the Japan Society for Promotion of Science Overseas fellowship.

## Author contributions

M.M., C.L., and A.I. conceived of and designed the project. M.M., C.L., and V.M. performed experiments. M.M., C.L., and V.M. analyzed and interpreted data. M.M. and A.I. wrote the manuscript and all authors reviewed and provided feedback on the manuscript. Authors from the Yale SARS-CoV-2 Genomic Surveillance Initiative contributed to sample screening, sample processing, SARS-CoV-2 genome sequencing, and data analysis.

## Data availability

All study data are included in the main text or Supplementary materials.

Yale SARS-CoV-2 Genome Surveillance Initiative members: Nicholas Chen, Mallery Breban, Anne M Hahn, Kien Pham, Tobias R Koch, Chrispin Chaguza, Irina Tikhonova, Christopher Castaldi, Shrikant Mane, Bony De Kumar, David Ferguson, Nicholas Kerantzas, David Peaper, Marie L Landry, Wade Schulz, Chantal BF Vogels, and Nathan D Grubaugh

## Supplementary figure legends

**Figure S1.**
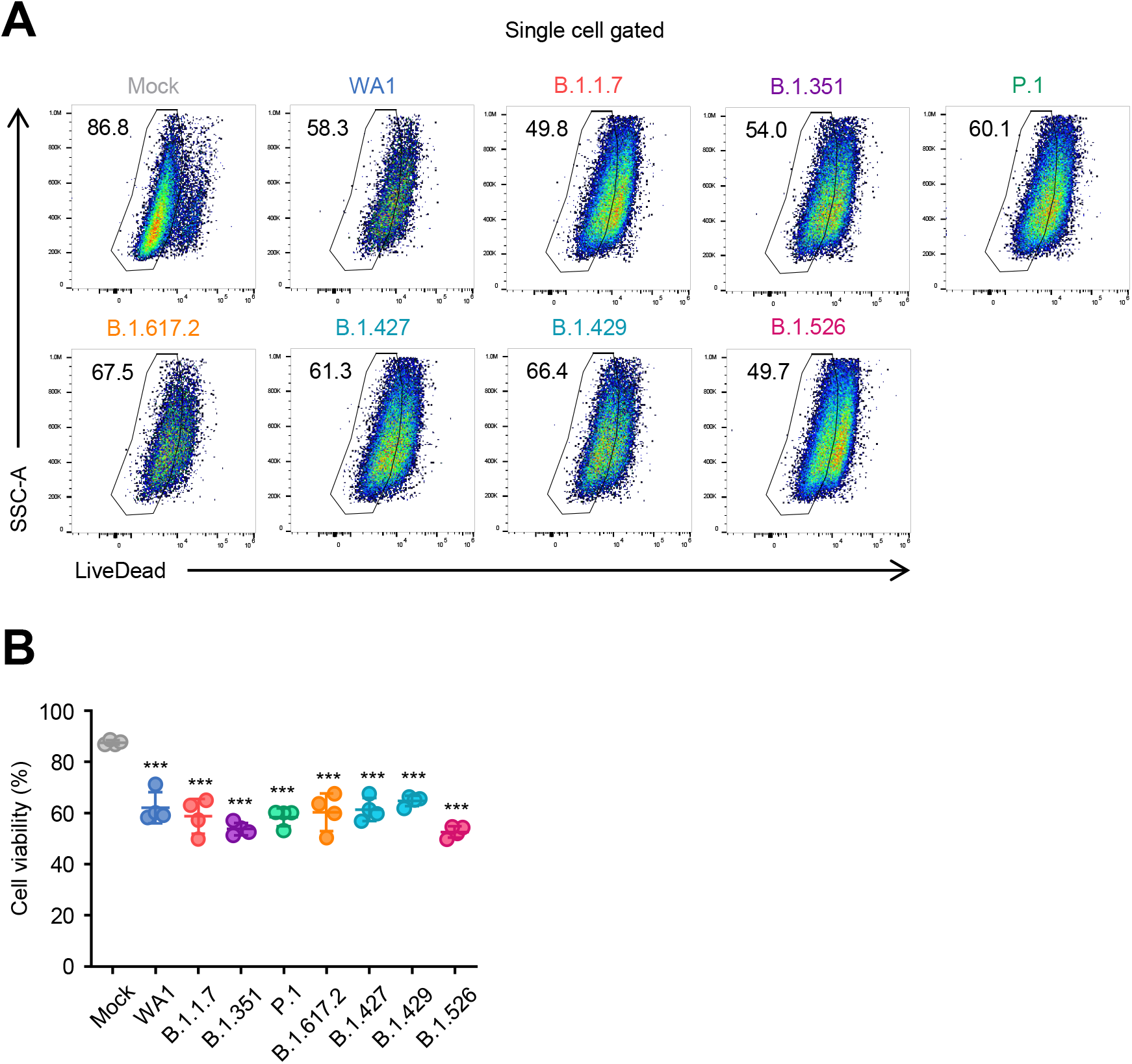
Cell viability during infection with SARS-CoV-2 variants. (A-B) Calu-3 cells were infected with SARS-CoV-2 variants at MOI 0.3 for 40h, and the cell viability was analyzed by FACS. (A) Representative FACS plot and (B) statistics are shown. Data are mean ± s.d. Data are representative of three independent experiments. ***, p< 0.001

**Figure S2.**
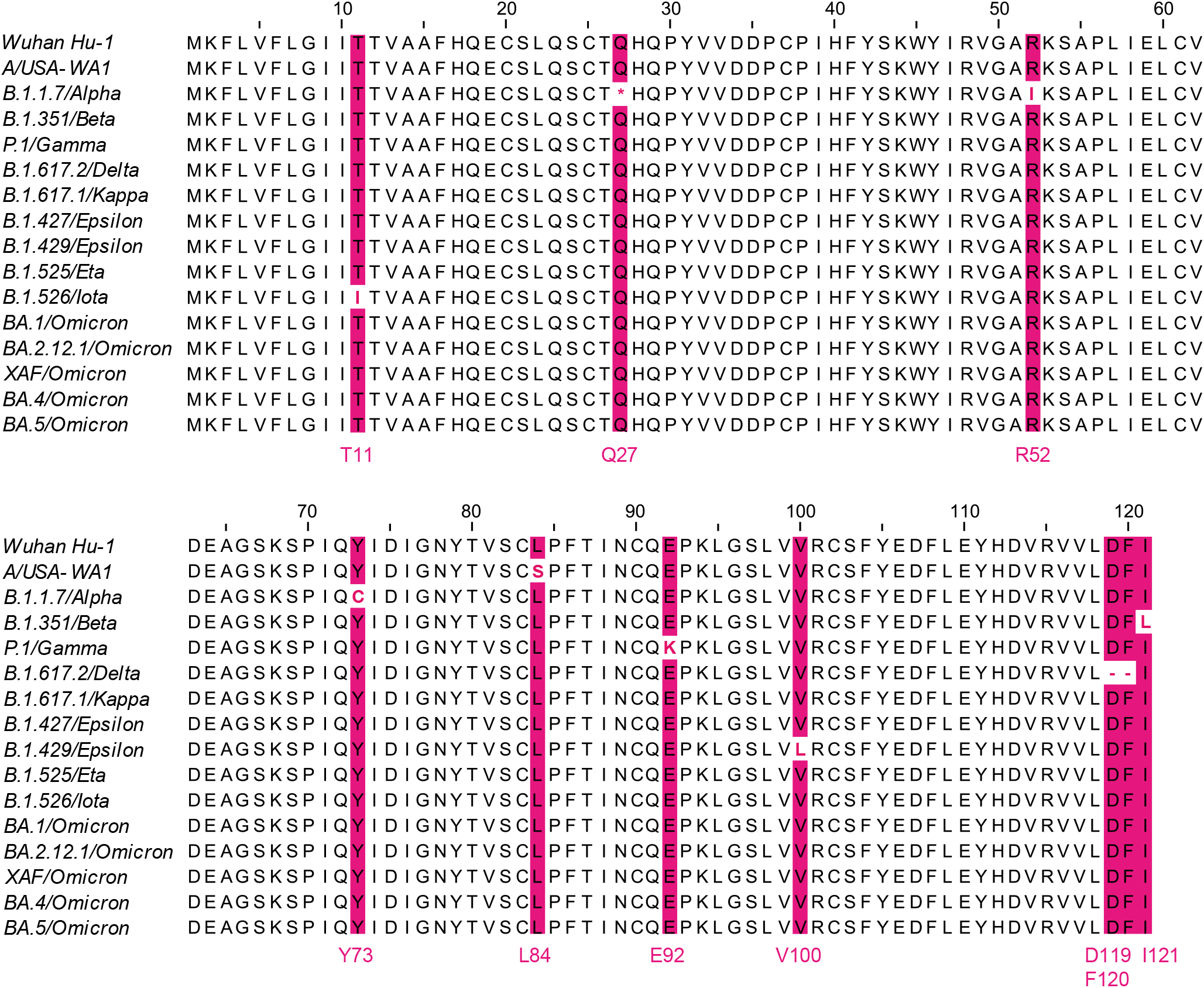
Comprehensive screening of mutations in ORF8 in variants of concern/interest. Multiple sequence alignment of ORF8 protein from SARS-CoV-2 variants. Amino acid residues where mutations are noted are colored in magenta.s.

**Figure S3.**
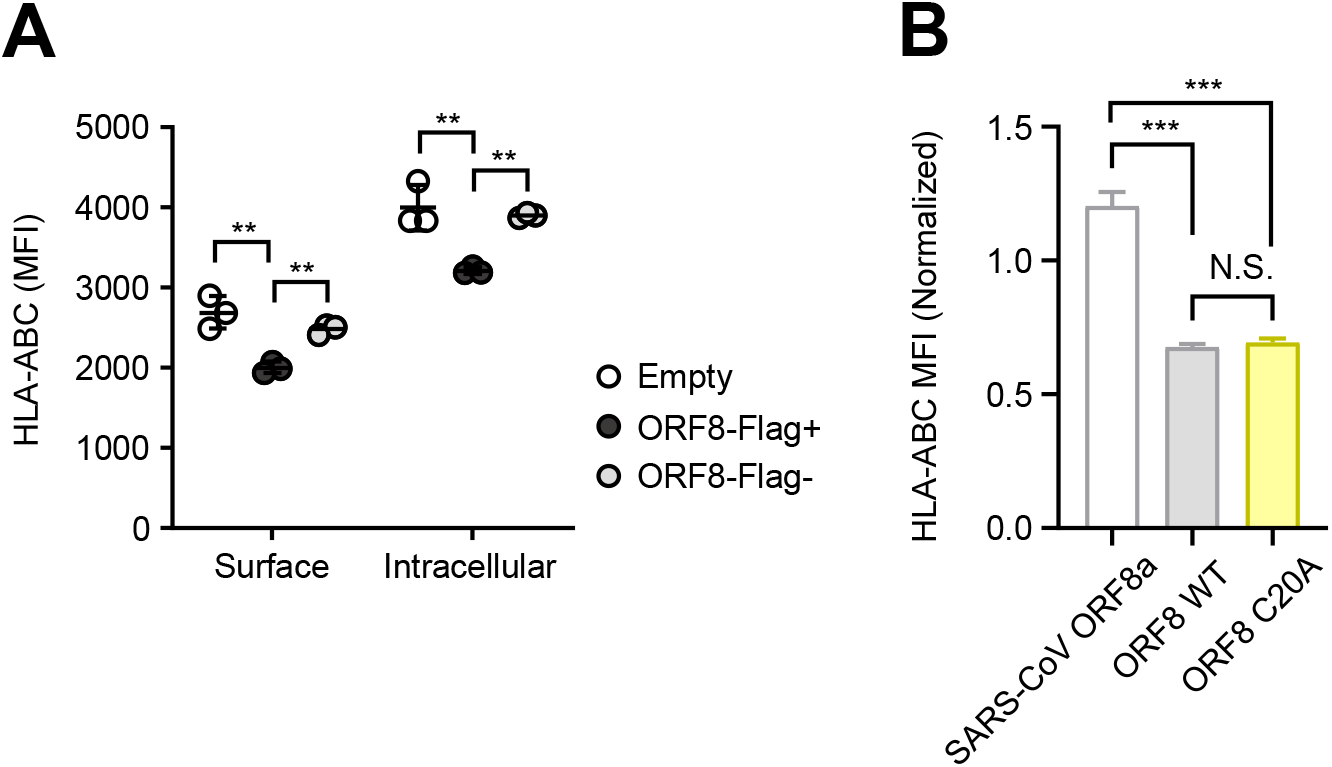
MHC-I evasion by SARS-CoV-2 ORF8. (A) HEK293T cells were transfected with plasmid encoding C-terminally Flag-tagged SARS-CoV-2 ORF8 WT or empty vector. Forty-eight hours after transfection, cells were collected and analyzed for the surface and intracellular HLA-ABC expression. (B) HEK293T cells were transfected with plasmids encoding C-terminally Flag-tagged SARS-CoV ORF8a, SARS-CoV-2 ORF8 WT, or SARS-CoV-2 ORF8 C20A mutant. Forty-eight hours after transfection, cells were collected and analyzed for the cell surface HLA-ABC expression. Data are shown as the ratio of MFI in Flag+ cells to Flag-cells (n=3). Data are mean ± s.d. Data are representative of two to three independent experiments. **, p< 0.01; ***, p< 0.001

